# Thermal stress responses of *Sodalis glossinidius*, an indigenous bacterial symbiont of hematophagous tsetse flies

**DOI:** 10.1101/638494

**Authors:** Jose Santinni Roma, Shaina D’Souza, Patrick J. Somers, Leah F. Cabo, Ruhan Farsin, Serap Aksoy, Laura J. Runyen-Janecky, Brian L. Weiss

**Author notes:** Corresponding authors: Laura Runyen-Janecky,; Brian L. Weiss.

## Abstract

Tsetse flies (Diptera: Glossinidae) house a taxonomically diverse microbiota that includes environmentally acquired bacteria, maternally transmitted symbiotic bacteria, and pathogenic African trypanosomes. *Sodalis glossinidius*, which is a facultative symbiont that resides intra and extracellularly within multiple tsetse tissues, has been implicated as a mediator of trypanosome infection establishment in the fly’s gut. Tsetse’s gut-associated population of *Sodalis* are subjected to marked temperature fluctuations each time their ectothermic fly host imbibes vertebrate blood. The molecular mechanisms that *Sodalis* employs to deal with this heat stress are unknown. In this study, we examined the thermal tolerance and heat shock response of *Sodalis*. When grown on BHI agar plates, the bacterium exhibited the most prolific growth at 25°C, and did not grow at temperatures above 30°C. Growth on BHI agar plates at 31°C was dependent on either the addition of blood to the agar or reduction in oxygen levels. *Sodalis* was viable in liquid cultures for 24 hours at 30°C, but began to die upon further exposure. The rate of death increased with increased temperature. Similarly, *Sodalis* was able to survive for 48 hours within tsetse flies housed at 30°C, while a higher temperature (37°C) was lethal. *Sodalis’* genome contains homologues of the heat shock chaperone protein-encoding genes *dnaK*, *dnaJ*, and *grpE*, and their expression was up-regulated in thermally stressed *Sodalis*, both *in vitro* and *in vivo* within tsetse flies. Arrested growth of *E. coli dnaK*, *dnaJ*, or *grpE* mutants under thermal stress was reversed when the cells were transformed with a low copy plasmid that encoded the *Sodalis* homologues of these genes. The information contained in this study provides insight into how arthropod vector enteric commensals, many of which mediate their host’s ability to transmit pathogens, mitigate heat shock associated with the ingestion of a blood meal.

**AUTHOR SUMMARY:** Microorganisms associated with insects must cope with fluctuating temperatures. Because symbiotic bacteria influence the biology of their host, how they respond to temperature changes will have an impact on the host and other microorganisms in the host. The tsetse fly and its symbionts represent an important model system for studying thermal tolerance because the fly feeds exclusively on vertebrate blood and is thus exposed to dramatic temperature shifts. Tsetse flies house a microbial community that can consist of symbiotic and environmentally acquired bacteria, viruses, and parasitic African trypanosomes. This work, which makes use of tsetse’s commensal symbiont, *Sodalis glossinidius*, is significance because it represents the only examination of thermal tolerance mechanisms in a bacterium that resides indigenously within an arthropod disease vector. A better understanding of the biology of thermal tolerance in *Sodalis* provides insight into thermal stress survival in other insect symbionts and may yield information to help control vector-borne disease.

## INTRODUCTION

Tsetse flies (Order: Diptera) house a microbial community that can consist of symbiotic and environmentally acquired bacteria, viruses, and parasitic trypanosomes. Among these are the primary endosymbiont *Wigglesworthia glossinidia*, a secondary symbiont *Sodalis glossinidius*, parasitic *Wolbachia sp.* (reviewed in (1, 2)) and *Spiroplasma* (3). *Sodalis* (order: Enterobacteriaceae) resides intra- and extracellularly within the fly’s midgut, hemolymph, milk and salivary glands, muscle, and fat body tissues (4–7). Both *Sodalis* and *Wigglesworthia* are passed vertically to tsetse progeny via maternal milk gland secretions (8, 9). Although the population dynamics of *Sodalis* in laboratory reared and field-captured tsetse flies has been well-documented (10–12), the functional relevance of this secondary symbiont to the fly’s physiology is currently unclear. *Sodalis* likely provides some benefit to tsetse, as flies exhibit a reduced lifespan when *Sodalis* is selectively eliminated via treatment with antibiotics (13). Additionally, *Sodalis* may modulate tsetse’s susceptibility to infection with parasitic African trypanosomes (*Trypanosoma brucei sp.*) (14), which are the etiological agents of human and animal African trypanosomiases. Specifically, this bacterium’s chitinolytic activity results in the accumulation of *N*-acetyl-D-glucosamine, which is a sugar that inhibits the activity of trypanocidal lectins (15). In support of this theory, several studies using field-captured tsetse have noted that the prevalence of trypanosome infections positively correlates with increased *Sodalis* density in the fly’s gut (16–19).

Because of the ectothermic nature of its tsetse host, *Sodalis* are likely exposed to a variety of temperatures in the fly’s natural niche. In particular, because tsetse is an obligate hematophagous insect, a rapid change in body temperature occurs during each feeding event that likely alters thermal stress physiology of the fly and its symbionts (20). In fact, temperature is one of the most important factors that controls bacterial growth and survival. Temperatures approaching and at the maximal temperature for a given bacterial species cause protein denaturation and membrane destabilization. Stabilization and refolding of denatured proteins via protein chaperones comprise mechanisms that bacteria use to combat thermal stress at elevated temperatures. One of the major chaperone systems is the ATP-dependent DnaK system, which also includes the co-chaperone DnaJ and the nucleotide exchange factor GrpE (reviewed in (21, 22)). DnaK homologues are distributed across all three branches of life. With the help of DnaJ, DnaK binds to unfolded proteins. ATP hydrolysis facilitates a conformational change in DnaK, which then surrounds the substrate protein and enables refolding. GrpE then facilitates ATP regeneration at the complex, which causes a conformational change that releases the refolded protein. In *E. coli* the expression of these three genes increases upon exposure to elevated temperatures via the alternative σ^32^ (σ^H^) sigma factor, which directs RNA polymerase to the *dnaK/dnaJ* and *grpE* promoters (23, 24).

The functional role of DnaK as it relates to thermal stress has been studied using a select number of model bacterial species. As such, this topic is understudied in symbionts that reside within ectothermic animal hosts. With respect to the DnaK/DnaJ chaperone system, Brooks *et al.* (25) showed that the system is required for *Vibrio fischeri* colonization of its *Euprymna scolopes* squid host via regulation of proper biofim formation. Manipulating genes of bacteria that are symbionts of insects, or that are vectored by insects, is technically challenging. As such, functional characterization of symbiont DnaK has heretofore been performed by ectopically expressing corresponding genes in heterologous bacteria. In these studies, the *dnaK* genes from *Buchnera aphidicola*, an aphid symbiont, and *Borrelia bordoferii*, which is vectored by ticks, showed partial to no complementation of the thermal sensitive phenotype in *E. coli dnaK* mutants (26, 27). Additionally, a second chaperone, GroEL, is one of the most highly expressed proteins in many insect bacterial symbionts, including *Sodalis* (28, 29).

The molecular mechanisms that underlie *Sodalis’* ability to reside successfully within the thermally fluctuating tsetse midgut environment are currently unknown. In this study we investigate *Sodalis’* thermal tolerance profile and the functionality of the bacterium’s DnaK/DnaJ/GrpE chaperone system in response to thermal stress.

## METHODS

### Bacterial strains, plasmids, and growth conditions

The bacterial strains and plasmids used in this study are listed in Table S1. *E. coli* strains were grown in Luria-Bertani Broth (LB) or on Luria-Bertani Agar (L Agar) plates. Liquid cultures were incubated at 37°C with 200 rpm aeration. *Sodalis glossinidius* were grown at 25°C and 10% CO_2_ on Brain Heart Infusion (BHI) Agar both with or without 10% horse blood (BHIB) (Haemostat Laboratories, Dixon, CA). The initial primary *Sodalis* culture used in these experiments (SOD^F^) was established by washing two week old *G. morsitans* pupae consecutively in 40% EtOH, 30% EtOH and sterile BHI media for 30 minutes per solution. Sterilized pupae were then homogenized in 100 μl of fresh BHI and plated on BBHI plates without antibiotics. Liquid *Sodalis* cultures were started by inoculating colonies into liquid BHI in petri dishes and incubated without aeration in a 10% CO_2_ microaerophilic environment. New cultures of *Sodalis* were typically inoculated at optical densities at 600 nm (OD_600_) of approximately 0.08. Antibiotics were used for *E. coli* at the following concentrations: carbenicillin (carb) 125 μg/ml, ampicillin (amp) 50 μg/ml, chloramphenicol (cam) 30 μg/ml, and kanamycin (kan) 50 μg/ml.

### Insect maintenance

*Glossina morsitans morsitans* were maintained in Yale’s insectary at 25°C with 60-65% relative humidity. All flies received defibrinated bovine blood (Hemostat Laboratories) every 48 hours through an artificial membrane feeding system (30).

### Plasmid constructions

Plasmids were isolated using the QIAprep Spin Miniprep Kit (Qiagen, Valencia, CA). DNA fragments and enzyme reactions were purified with QIAquick gel extraction kit (Qiagen). All ligations were done with T4 DNA Ligase (Promega) and transformed into *E. coli* DH5α. The sequences of the PCR primers for cloning are listed in Table S2, and all PCRs for cloning were done using PfuTurbo (Agilent, Santa Clara, CA). To construct pJR1 and pJS2, DNA fragments containing *dnaK* were PCR amplified from *Sodalis* or from *E. coli* (with primers UR423 and UR424 or UR518 and UR519, respectively), digested with XbaI and XhoI, and ligated to pWKS30 digested with the same enzymes. To construct pJR5, a modified procedure from the Quickchange II Site Directed Mutagenesis Kit (Agilent Technologies, Santa Clara, CA) was used. A DNA fragment containing *dnaJ* and ends that match the pRJ1 plasmid was PCR amplified from *Sodalis* with primers UR530 and UR531. This PCR fragment was then annealed to pJR1 and used as the primers to synthesize a larger plasmid containing both *Sodalis dnaK* and *dnaJ*. The template pJR1 plasmid was then degraded with addition of DpnI, and the reaction mixture was transformed into *E. coli* NEB 5-alpha to recover pJR5. This plasmid was sequenced to verify that the DNA sequence of *dnaKJ* was the same as the chromosomal sequence. To construct pSD2, pJR5 was cut with MscI and a 1 kb internal segment of *dnaK* was deleted. To construct pRF2, a DNA fragment containing *grpE* was PCR amplified from *Sodalis* with primers UR455 and UR456 digested with XbaI and PstI, and ligated to pWKS30 digested with the same enzymes.

### Quantitation of *Sodalis* thermal tolerance gene expression

For measuring *in vitro* gene expression, *Sodalis* cultures were grown in BHI from a starting OD_600_ of 0.08 for 1 day and then thermally stressed as follows: Cultures (1 ml) were transferred to microfuge tubes and placed in water baths at 25°C or 30°C for 15 minutes. 250 μl of RNA stabilizing reagent (95% acidic phenol/5% ethanol) was added to each sample to stabilize the RNA after incubation. The RNA was isolated using the RNeasy Mini Kit Procedure (Qiagen), and the isolated RNA was treated with DNase I until the samples were DNA-free (Qiagen).

For measuring *in vivo* gene expression, tsetse (3 biological replicates, *n*=5 flies per replicate) were housed at 37°C for 1 hour. Controls (3 biological replicates, *n*=5 flies per replicate) were maintained at 24°C. Following exposure to thermal stress, midguts were rapidly dissected from each fly and transferred to liquid nitrogen for stabilization. Samples (pools of 5 guts each) were homogenized on ice in 240 μl Tri-Reagent (Zymo Research, Irvine, CA) using a motorized homogenizer and then debris was removed by brief centrifugation. Total RNA (including symbiont RNA) was then isolated using the Direct-zol RNA Kit (Zymo Research), and the isolated RNA was treated with DNase I (Qiagen) until the samples were DNA-free.

cDNA was generated from 200 ng total RNA (*in vitro*) and 880 ng total RNA (*in vivo*) using Superscript III and random hexamers (Invitrogen, Carlsbad, CA). Quantitative PCR was performed on the cDNA samples using primers UR545 and UR546 for *dnaK*, UR556 and UR557 for *dnaJ*, and UR558 and UR559 for *grpE*. Primers QrplB1F and QrplB1R, which amplify the constitutively expressed gene *rplB* (which had been previously verified (31, 32)), were used as a control to ensure equal amounts of cDNA in each sample.

### Thermal stress assays for *Sodalis in vitro*

For growth assays, *Sodalis* were inoculated in BHI at an OD_600_ of 0.08, grown for 2 days at 25°C, re-normalized to an OD of 0.02 (~10^7^ bacteria per ml), and serially diluted in BHI broth. 10 μl (or 100 μl) of each serial dilution was spotted (or spread) on agar plates [BHI, BHIB or BHI supplemented with 0.6-10% horse serum (Hemostat)]. Plates were incubated in a 10% CO_2_ incubator or in CampyPak microaerobic pouches (Becton, Dickinson and Company, Franklin Lakes, NJ) at the various temperature as indicated in the figure and table legends. Growth under each condition was assessed by scoring for the presence or absence of colonies after 7 days.

For survival assays, *Sodalis* were inoculated in BHI at an OD_600_ of 0.08, grown for 2 days at 25°C, diluted to an OD_600_ of 0.1 in BHI broth, and aliquoted into samples that were incubated at 25°C, 30°C, 32°C, or 37°C for 3 days. Viable surviving bacteria were quantified each day by serially diluting the samples and plating on BHI agar plates and incubating the plates at 25°C. The number of colonies was counted seven days later, total number of viable cells at each temperature and time point were calculated by multiplying the number of colonies by the dilution factor and dividing by the amount of sample plated.

### Thermal stress assays for *Sodalis in vivo*

*Sodalis* residing within tsetse were thermally stressed by holding flies at 30°C or 37°C in an incubator for 48 hours. Subsequently, midguts microscopically excised from these flies, and control flies maintained at 25°C, were homogenized in 0.85% NaCl, serially diluted and plated on BHIB agar (10) Colony forming units per plate were counted manually.

### Thermal stress assays for *E. coli* thermal stress mutants expressing *Sodalis* genes

Overnight cultures of the *E. coli* strains carrying plasmids with *Sodalis* genes were grown in LB containing carb and the antibiotic marking the chromosomal mutations (cam for MC4100Δ*dnaK* and kan for JW0014-1) at 30°C. For assessment of growth on agar plates, the overnight cultures were serially diluted 1:10 six times in LB and 10 μL each dilution were spotted onto L agar plates containing carb. Plates were placed at 30°C (non-stress temperature) and 45+1°C (thermal stress temperature) overnight and growth was assessed the following day. For assessment of growth in liquid, the overnight cultures were subcultured 1:100 in LB containing carb and incubated at 30°C until midlog stage was reached. Then, each culture was diluted to an OD_600_ of 0.06 in LB containing carb and incubated at either 30°C or 46°C. Growth was measured via optical density at 600 nm.

### Statistical analyses

All statistical analyses were carried out using GraphPad Prism (v.8). All statistical tests used, and statistical significance between treatments, and treatments and controls, are indicated on the figures or in their corresponding legends. All samples sizes are provided in corresponding figure legends or are indicated graphically as points on dot plots. Biological replication implies distinct aliquots of cultured *Sodalis*, and distinct groups of flies collected on different days, were used for all experiments.

## RESULTS

### Establishing the thermal range for *Sodalis* growth *in vitro* on un-supplemented BHI agar

To determine maximal temperature for *Sodalis* growth, *Sodalis* cultures were serially diluted and spotted on BHI agar plates, which were then incubated at various temperatures for 7 days. All of the spotted *Sodalis* dilutions (ranging from 10^−1^ to 10^−4^) formed colonies on BHI agar at temperatures up to 29°C. At 30°C on BHI, *Sodalis* was unable to form colonies at the highest two dilutions (10^−3^ and 10^−4^) and did not form any colonies at ≥ 31°C (Table 1, column 1).

**Table 1.** Thermal growth range for *Sodalis* on agar plates.

| incubation<br>temperature (°C) | Growth of bacteria<sup>a</sup> |  |  |  |
| --- | --- | --- | --- | --- |
|  | in CO<sub>2</sub> incubator |  | in CampyPak |  |
|  | BHI | BHI + blood | BHI | BHI + blood |
| 25 | + | + | + | + |
| 29 | + | + | + | + |
| 30 | ± ($10^{-2}$) | + | + | + |
| 31 | | ± ($10^{-2}$) | ± ($10^{-2}$) | ± ($10^{-3}$) |
| 32 | | | Haze at $10^{-1}$ | ± ($10^{-2}$) |
| 33 | | | | Haze at $10^{-1}$ |

<sup>a</sup>Two day old cultures of *Sodalis* were normalized to an OD of 0.02 ( $\sim 10^7$ bacteria per ml) and serially diluted 1:10 four times. 10 $\mu\text{l}$ of each serial dilution was spotted on agar plates and incubated as indicated. The data shown are representative of at least three independent experiments. +, growth at all dilutions; ± , growth only to the dilution indicated in parenthesis. Black shaded boxes = no visible growth.

### Presence of blood in the agar or growth in microaerobic environments extends *Sodalis* thermal tolerance

Tsetse’s gut is likely hypoxic, and periodically replete with blood. Thus, to more closely mimic gut conditions and determine how they influence *Sodalis* thermal stress survival, we tested the bacterium’s thermal tolerance when grown on BHI agar plates supplemented with blood and/or in a reduced oxygen environment (through the use of CampyPak microaerobic pouches, which reduces environmental oxygen levels to 5-15%). We found that either condition enabled *Sodalis* to form colonies at the highest dilution (10^−4^) at 30°C (Table 1). However, as the temperature increased, the ability of the bacteria to form colonies at all dilutions was diminished in these supplemented growth conditions. At 32°C, no colonies were present on the blood-supplemented plate incubated aerobically, and only a hazy film of growth formed at 10^−1^ upon CampyPak-incubation. The plates that had both blood-supplementation and CampyPak-incubation showed colonies at 10^−2^ dilutions at 32°C, but just a haze of growth at 33°C (Table 1).

Growth stimulation due to blood supplementation of agar could be due to either a component of the erythrocytes or the serum fraction. To determine which contained the stimulating factor(s), we added either purified erythrocytes or serum to the BHI agar plates. Only the agar supplemented with the erythrocytes supported growth (Table 2), suggesting that some component of this cell type facilitates *Sodalis* growth under stressed conditions.

**Table 2.**
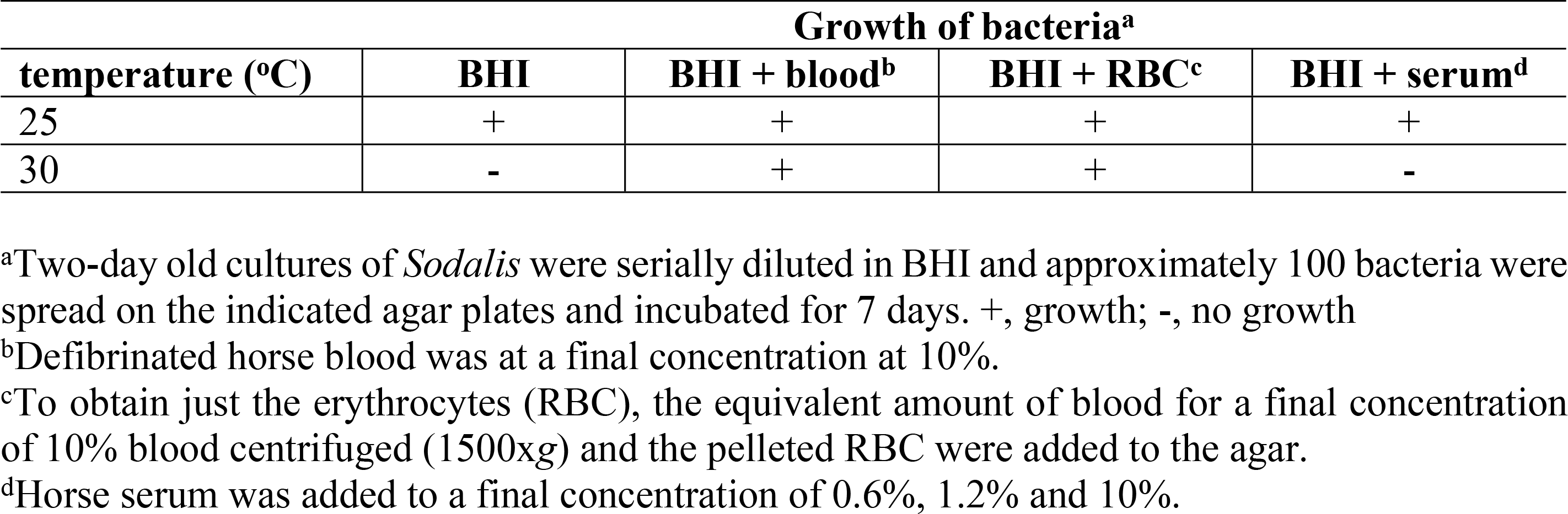
Blood stimulation of *Sodalis* growth.

| temperature (°C) | Growth of bacteria<sup>a</sup> |  |  |  |
| --- | --- | --- | --- | --- |
|  | BHI | BHI + blood<sup>b</sup> | BHI + RBC<sup>c</sup> | BHI + serum<sup>d</sup> |
| 25 | + | + | + | + |
| 30 | - | + | + | - |

<sup>a</sup>Two-day old cultures of *Sodalis* were serially diluted in BHI and approximately 100 bacteria were spread on the indicated agar plates and incubated for 7 days. +, growth; -, no growth
<sup>b</sup>Defibrinated horse blood was at a final concentration at 10%.
<sup>c</sup>To obtain just the erythrocytes (RBC), the equivalent amount of blood for a final concentration of 10% blood centrifuged (1500xg) and the pelleted RBC were added to the agar.
<sup>d</sup>Horse serum was added to a final concentration of 0.6%, 1.2% and 10%.

### Exposure to temperatures above 30°C is lethal to *Sodalis in vitro* and *in vivo*

The lack of *Sodalis* growth above 30°C on BHI agar could be because this temperature is either bacteriostatic or bactericidal (bacteria are alive but not growing, versus bacteria are dead). To distinguish between these two possibilities, we exposed *Sodalis* cultured in BHI broth to temperatures that were non-permissive for optimal growth (≥ 30°C) and then plated the bacteria on agar plates at a permissive temperature to quantify the number of cells that remained viable in the BHI broth culture. Within 24 hours of exposure to the non-permissive temperatures, all samples incubated *above* 30°C had less than 40% survival; by 72 hours, only 0.4% of the cells were recovered from samples exposed to 32°C, and no bacteria could be recovered from the samples at 37°C (Fig. 1A). The samples incubated at 30°C were able to survive as well as the samples at 25°C for the first 24 hours. However, only 33% and 12% of the cells survived for 48 and 72 hours, respectively.

**Fig. 1:**
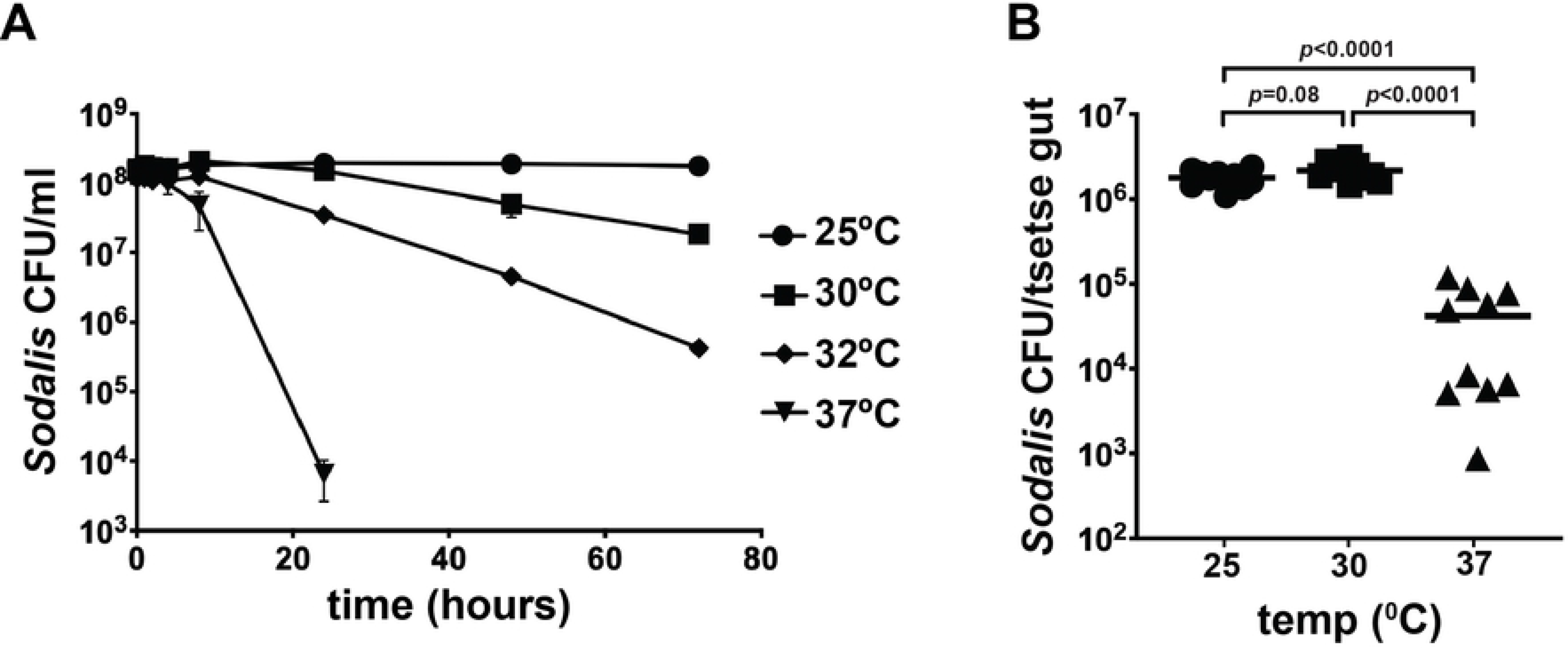
*Sodalis* survival at non-permissive growth temperatures. (A) Two-day old *Sodalis* cultures were diluted to an OD_600_ of 0.1 and incubated at 25°, 30°C, 32°C and 37°C for 3 days. Viable surviving bacteria were quantified by plating on BHI agar plates at the indicated time points and incubating the plates at 25°C. The number of colonies was counted seven days later and used to determine the total number of viable cells at each temperature and time point. Each timepoint represents the average of three trials, ± standard deviation. (B) Tsetse flies were reared at 25°C, 30°C and 37°C for 48 hours, after which midguts were excised, homogenized in 0.85% NaCl and plated on BBHI/agar. *Sodalis* density per midgut was determined by manually counting colonies. Each data point on the graph represents one midgut (*n*=10 per treatment), and statistical significance was determined using a one-way ANOVA followed by Tukey’s HSD post-hoc analysis.

We next investigated the thermal tolerance of *Sodalis* that reside indigenously within tsetse by exposing wild-type flies to elevated temperatures for 48 hours. Similar numbers of *Sodalis* were recovered from control (1.8×10^6^±1.3×10^5^ CFU) and treatment (2.2×10^6^±1.8×10^5^ CFU) tsetse housed at 25°C and 30°C, respectively. Conversely, tsetse housed at 37°C harbored 4.2×10^4^±1.3×10^4^ *Sodalis* CFU, which represents a 98% reduction in bacterial density compared to controls (Fig 1B).

### *Sodalis* contains DnaK, DnaJ, and GrpE homologues

The DnaK/DnaJ/GrpE chaperone system enables bacterial cells to respond to elevated temperature (reviewed in (22)). *Sodalis’s* genome encodes proteins annotated as DnaK, DnaJ, and GrpE (33). The *dnaK* and *dnaJ* genes are tandemly organized on the chromosome, with a 117 bp intergenic region between them, while *grpE* is located 2 Mbp away. Well-conserved σ^32^ binding sites are located 124 and 47 bp 5’ of the *dnaK* and *grpE* start codons, respectively (Fig. S1A and S1B). No σ^32^ or σ^70^ binding sites are present in the intergenic region between *dnaK* and *dnaJ*, but RNAs containing this sequence are capable of folding into an extended hairpin (Fig. S1C).

*Sodalis* is less able to survive high temperatures than is closely-related, free-living *E. coli*. These differential phenotypes prompted us to compare the amino acid sequences of the *Sodalis* and *E. coli* chaperone proteins. *Sodalis* DnaK and DnaJ are 89% identical to their *E. coli* homologues, while *Sodalis* and *E. coli* GrpE exhibit 65% identity. Additionally, these *Sodalis* proteins have retained the conserved residues implicated in conferring thermal tolerance in *E. coli* (Fig. S2). We also examined the genomes of other insect bacterial symbionts for *Sodalis* DnaK homologues by performing a BLAST search with *Sodalis* DnaK. We found high levels of DnaK conservation with *Sodalis* DnaK in a wide variety of insect symbionts, including the primary symbiont of tsetse *Wigglesworthia glossinidia* (Fig. S3, Table S3). Additionally, these proteins also retain conserved residues that confer thermal tolerance in *E. coli*. Our findings suggest that distinct bacterial taxa (e.g., *E. coli* and *Sodalis*) may exhibit very different thresholds of thermal stress despite the highly conserved nature of thermal heat shock genes. Furthermore, elevated expression of these genes occurs at different temperatures, and other proteins may be involved in modulating thermal stress, in different bacteria.

### *dnaK* transcription increases at elevated temperatures *in vitro and in vivo*

In *E. coli* and many other bacteria, temperatures above the optimal growth temperature induce expression of *dnaK (34)*. Thus, we hypothesized that *Sodalis dnaK* expression would similarly increase at temperatures above 25°C. To test this hypothesis, we isolated total RNA from *Sodalis* exposed briefly to 25°C and 30°C temperatures *in vitro* and then used RT-qPCR to measure *dnaK*, *dnaJ*, and *grpE* expression levels. *Sodalis dnaK* expression increased 6-fold when the bacterium was exposed to 30°C, as compared to 25°C (Fig. 2). Likewise, *dnaJ* and *grpE* expression was 9-fold and 7-fold induced, respectively, at elevated temperatures (Fig. 2).

**Fig. 2.**
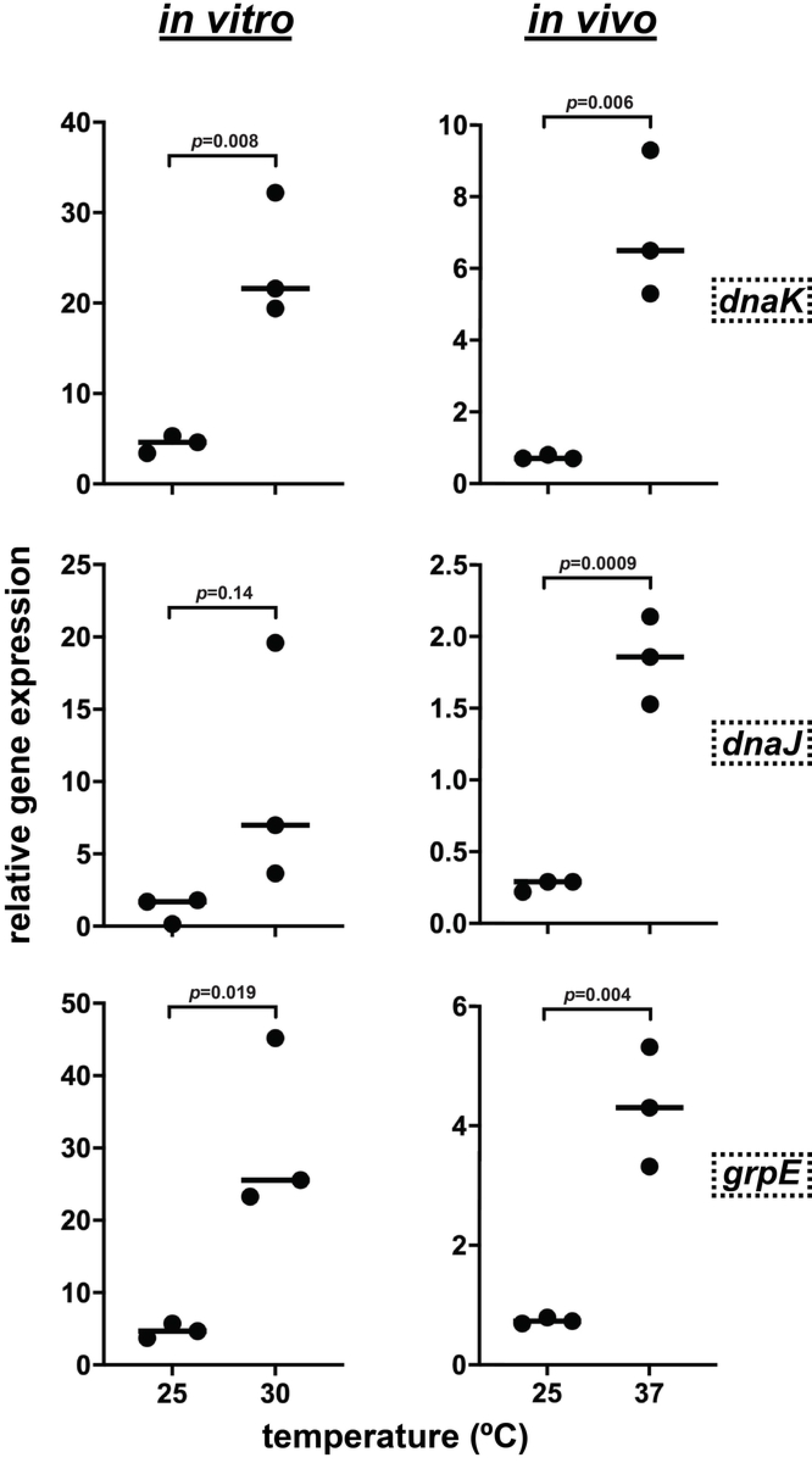
*Sodalis* chaperone gene expression increases with thermal stress. For *in vitro* experiments (left panels) *Sodalis* were grown in BHI at 25°C for 1-2 days. Subsequently, the culture was split into two samples that were incubated at either 25°C or 30°C for 15 min. RNA was then isolated from each sample and used to generate cDNAs, which were amplified by quantitative real time PCR using *dnaK*, *dnaJ* and *grpE* specific primers. The amount of *dnaK, dnaJ*, and *grpE* expression was normalized to the housekeeping gene *rplB* by dividing the relative amounts of each cDNA by the relative amounts of *rplB* cDNA in each sample. For *in vivo* experiments (right panels) tsetse flies were housed at either 25°C or 37°C for 1 hour. Total RNA was extracted from the guts of these tsetse flies and converted into cDNA. *Sodalis dnaK* (panel A), *dnaJ* (panel B), and *grpE* (panel C) were then amplified from these cDNAs using quantitative real time PCR. The amount of *dnaK, dnaJ*, and *grpE* expression was normalized to the housekeeping gene *rplB* by dividing the relative amounts of each cDNA by the relative amounts of *rplB* cDNA in each sample. On all graphs each data point represents one biological replicate (*n*=5 fly guts per replicate), and the line indicates the median of the replicates. Statistical significance for all experiments was determined using unpaired t-tests.

We performed a similar experiment with symbiont-carrying tsetse flies housed at 25°C versus a cohort maintained at 37°C for 1 h. This latter temperature shift mimics that encountered by enteric *Sodalis* when tsetse imbibes a vertebrate blood meal. Expression of all three genes (*dnaK, dnaJ*, and *grpE*) increased 6-10 fold when tsetse flies were exposed to 37°C (Fig. 2), indicating that *Sodalis*’ thermal stress tolerance system is active in the bacterium’s native environment.

### *Sodalis* DnaK, DnaJ, and GrpE mediate thermal tolerance in a heterologous *E. coli* host

To test whether the *Sodalis* DnaK, DnaJ, and GrpE proteins are functional chaperones, we expressed these *Sodalis* genes on plasmids in several different *E. coli* strains that lack the respective homologues. *E. coli* MC4100ΔdnaK, JW0014-1, and DA16 lack functional DnaK, DnaJ, and GrpE, respectively, and these strains cannot grow at temperatures at or above the elevated temperature of 45°C. To quantify the ability of the *Sodalis* chaperone proteins to functionally replace their *E. coli* homologues, we spotted serially diluted cultures of the mutant *E. coli* strains that express *Sodalis* genes on plates that were then incubated at permissive and elevated temperatures. All strains grew at the permissive temperature of 30°C (Table S4). At 45°C, the *E. coli* mutant strains containing pWKS30, the vector control (empty plasmid), failed to grow. However, *E. coli* MC4100ΔdnaK expressing *Sodalis dnaK* survived as well as the parent strain at this elevated temperature (Fig. 3). Likewise, *E. coli* JW0014-1 (Δ*dnaJ*) and DA16 (Δ*grpE*) expressing *Sodalis dnaJ* or *Sodalis grpE*, respectively, survived elevated temperature as well as the parent strains (Fig. 3).

**Fig. 3.**
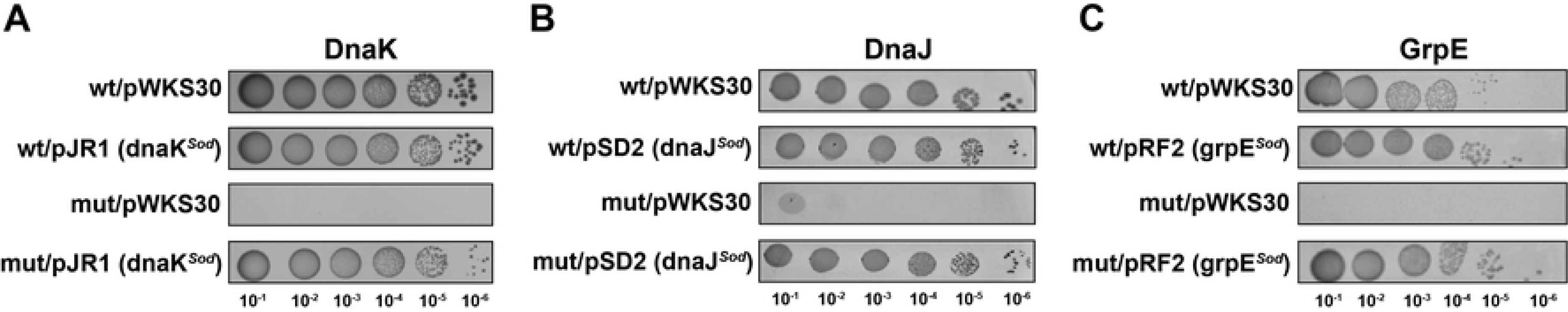
*Sodalis* chaperone genes facilitate *E. coli* survival at elevated temperatures. Overnight cultures of the indicated *E. coli* strains grown at 30°C were serial diluted (1:10) six times, and 10 μl of each dilution was spotted on L agar plates. The plates were incubated at 45±1°C for 18-24 hours. In all panels, experimental designations are indicated as the *E. coli* strain/introduced plasmid (containing the cloned *Sodalis* gene). All plasmids used are described in Table S1. Panel A, wt (wild-type *E. coli* strain MC4100), mut [mutant *E. coli* strain MC4100ΔdnaK (*ΔdnaK*)]. Panel B, wt (wild-type *E. coli* strain BW25113, mut [mutant *E. coli* strain JW0014-1 (*ΔdnaJ*)]. Panel C, wt wild-type *E. coli* strain DA15, mut [mutant *E. coli* strain DA16 (*grpE280*)]. The data shown are representative of at least three independent experiments.

We also examined the growth kinetics of *E. coli* strains expressing *Sodalis dnaK* or *dnakJ* by performing growth curves at 30°C and 46°C. All strains grew at the permissive temperatures of 30°C (Fig. 4), although the *dnaK* mutant containing the pWKS30 control plasmid grew slightly slower than the wildtype strain containing pWKS30. At 46°C, the *E. coli dnaK* mutant strain containing pWKS30 did not grow. *E. coli* MC4100ΔdnaK expressing *Sodalis dnaK* or *Sodalis dnaKJ* survived as well as the parent strain and control strains with *E. coli dnaK* (Fig. 4). Taken together, these data suggest that the *Sodalis* DnaKJ/GrpE chaperone system is sufficient for mediating heat shock survival in an *E. coli* heterologous host strain deficient in these functions.

**Fig. 4.**
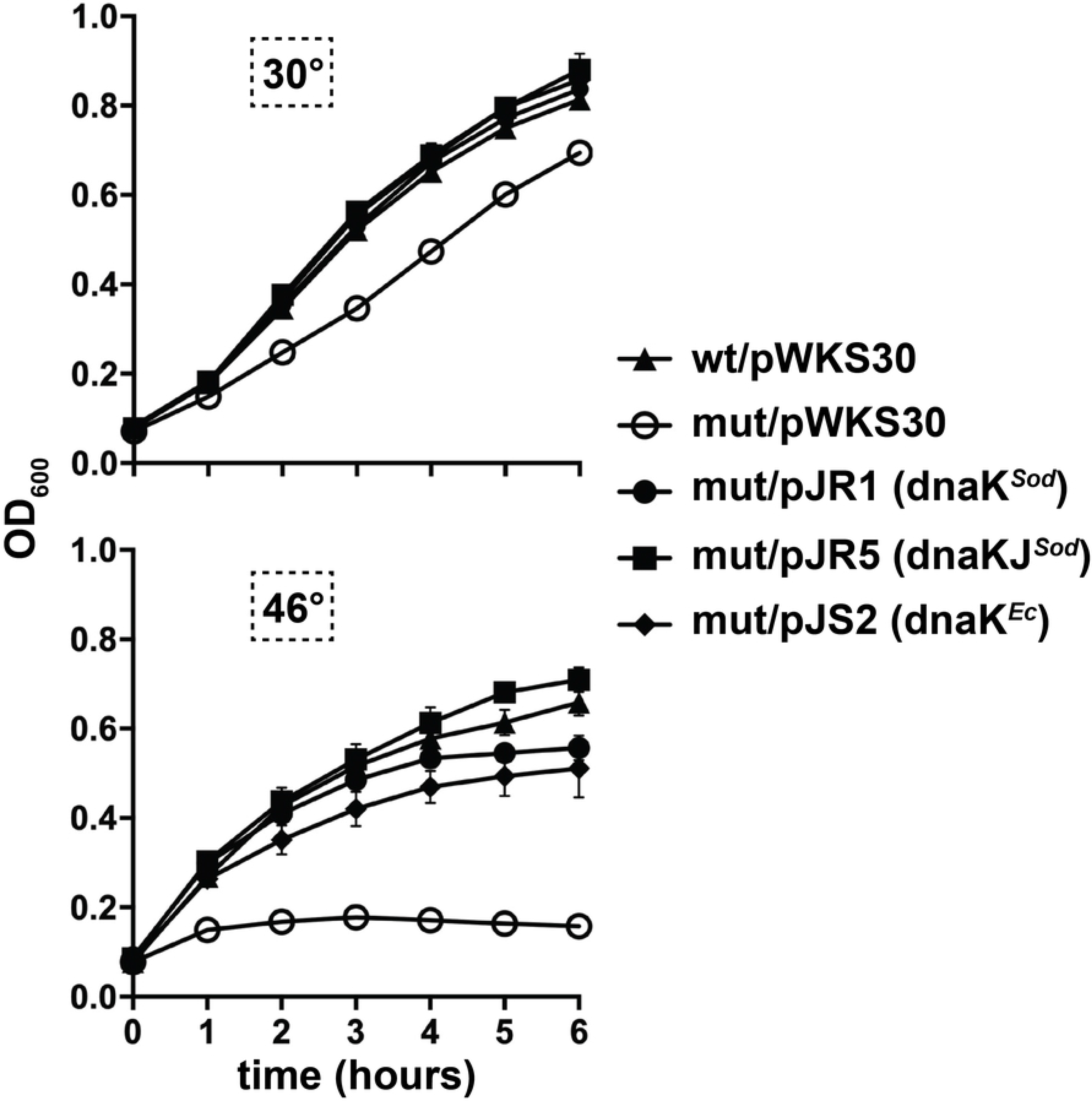
Growth kinetics of *E. coli dnaK* mutants containing *Sodalis dnaK* or *dnaK and dnaJ*. Mid-log phase cultures (37°C) of *E. coli* wt/pWKS30 or mutant *E. coli* [MC4100ΔdnaK (*ΔdnaK*)] containing either pWKS30, pJR1, pJR5, or pJS2 (all plasmids used are described in Table S1) were diluted to an OD_600_ of 0.06 in L broth containing carbenicillin and incubated at 30°C (top graph) or 46°C (bottom graph). Growth was measured via optical density at 600 nm. Each time point represents the mean of three individual experiments, ± the standard deviation.

## DISCUSSION

Ectothermic insects, and the bacterial symbionts that reside within them, are sensitive to temperature shifts in their environment. Bacteria that reside within the gut of hematophagous insects must also deal with a rapid increase in the temperature of their niche when their host consumes vertebrate blood. This study utilized the tsetse fly and its secondary symbiont, *Sodalis glossinidius*, to examine molecular mechanisms that may mediate thermal stress tolerance in an obligate blood feeding insect/bacterial symbiont model system. Our initial work examined the upper thermal limit for growth and survival of *Sodalis* maintained in culture and residing endogenously within tsetse. We found that *Sodalis* do not survive for extended periods of time when exposed to temperatures above 30°C in BHI media or within tsetse at 37°C. This was surprising because, outside of a lab setting, tsetse flies and their *Sodalis* symbionts reside in sub-Saharan Africa where temperatures are often well above 30°C. Several scenarios may explain this unexpected finding. First, tsetse are crepuscular and thus feed during cooler dawn and dusk periods of the day (35). Furthermore, although the environmental temperature routinely exceeds 30°C, tsetse seek cooler shade under these conditions (36). Thus, even in the wild, *Sodalis* within tsetse might not be exposed to temperatures above 30°C for long periods of time. Although the fly’s body temperature rises during and immediately after feeding to 36°C (20), it likely returns to ambient temperature well within *Sodalis’* window of survival. In fact, our results showed that in BHI broth, *Sodalis* can survive for limited periods of time at 30°C for 24 hours, 32°C for 8 hours, and 37°C for at least 2 hours. Second, the *Sodalis* used in this study were isolated from tsetse flies reared in a laboratory colony at 25°C for many years. *Sodalis* from these flies may have lost the ability to survive at higher temperatures, while recent environmental isolates of *Sodalis* may be more thermo-tolerant. Finally, environmental factors associated with the tsetse fly host (including other symbionts) may increase the upper thermal limit for *Sodalis* survival, and these factors may be absent from the tsetse colony and BHI growth media.

We discovered that modulating the composition and/or environment of agar plates on which *Sodalis* are cultured can restore growth at temperatures above 30°C. Specifically, either blood-supplementation of the agar, or growth within hypoxic CampyPaks (5-15% O_2_), increased the high temperature threshold at which *Sodalis* could grow. The stimulation of growth by addition of whole blood is likely due to a component of the erythrocytes, and not the serum, as addition of serum alone did not restore growth of *Sodalis* at 30°C. In addition to hemoglobin, erythrocytes contain high levels of catalase (37, 38), which detoxifies the oxidative stress molecule H_2_O_2_ that is generated during cellular metabolism. Thus, catalase-mediated mitigation of oxidative stress in the presence of blood/erythrocytes may allow *Sodalis*, which does not itself produce catalase (39), to divert more resources to surviving thermal stress situations. Consistent with this hypothesis are the observations that (a) incubation of *Sodalis* in CampyPaks, which reduce oxidative stress by generating a hypoxic environment, also increased the high temperature threshold at which *Sodalis* could grow, but addition of hemoglobin did not, and (b) thermal tolerance and oxidative stress are physiologically linked, as increased temperatures can result in an oxidative stress burden on bacteria cells (40, 41) (42). A similar phenomenon might be occurring in *Sodalis* in the fly. Specifically, tsetse’s gut is likely hypoxic, and this environment may increase the thermal limit for *Sodalis* within the fly. Consistent with this idea, we found that although *Sodalis* struggled to grow at 30°C on BHI agar, the bacteria grew fine in the tsetse fly gut at this temperature.

DnaKJ/GrpE chaperone systems that enable refolding of proteins during thermal stress have been found in all domains of life, suggesting that selection for maintenance of these systems is strong. An analysis of over 1200 genomes in 2012 showed that *dnaK* was found in all bacteria, with the exception of two thermophiles isolated from deep sea hydrothermal vents (43). More so, the function of DnaK chaperone systems is well characterized in several bacteria, the vast majority of which are pathogens. Comparatively, genomic conservation and physiological function systems is less well studied in symbionts that reside within ectothermic animal hosts. One exception is the aphid symbionts *Buchnera*, *Serratia symbiotica* and *Hamiltonella defensa*, the latter two of which facilitate their insect host’s survival in high temperature environments (51). Additionally, *dnaJ* and *grpE* are lost or truncated in a few highly reduced genomes from vertically transmitted endosymbionts that share ancient associations with cicadas, mealybugs, and psyllids (43). Our work herein shows that the *dnaKJ* locus is conserved between *Sodalis* and *E. coli*, including promoter regions and a potential hairpin in the intergenic region between *dnaK* and *dnaJ*. Additionally, at the protein level, *Sodalis* DnaK is 89% identical to its *E. coli* ortholog, and residues implicated in mediating thermal tolerance are conserved. Thus, the fact that *Sodalis* DnaK, DnaJ, and GrpE proteins can functionally replace the homologous *E. coli* proteins and promote growth at elevated temperature is not surprising. However, this finding of complementation is in contrast with DnaK from *Buchnera aphidicola*, *Borrelia burgdorferi* and *Vibrio harveyi*, which exhibit partial (*B. burgdorferi*, at 37°C but not 43°C) or no complementation phenotypes when expressed ectopically in *E. coli dnaK* mutants (27, 44). Our ability to complement was similar to complementation found with DnaK from *Pseudomonas syringae*, *Agrobacterium tumefaciens*, and *Brucella ovis* (45–47). No apparent correlation exists between evolutionary relatedness of the above-mentioned bacteria to *E. coli* and the ability of their DnaK proteins to complement an *E. coli dnaK* mutant.

For bacteria that experience rapid fluctuations in their environment, the DnaK chaperone system may be especially critical. Induction of *Sodalis dnaK* expression in response to elevated temperature suggests that the bacterium uses DnaK to tolerate heat shock. Retention of these systems in tsetse’s indigenous, enteric symbionts is indicative of their fundamental importance during times of thermal stress, such as that experienced when the fly host consumes vertebrate blood. In support of this theory, the highly reduced (~ 700 kB) genome of tsetse’s obligate mutualist, *Wigglesworthia*, also encodes a conserved and putatively functional *DnaK* gene (48) This bacterium resides intracellularly within tsetse’s bacteriome organ (49), which is attached to the fly’s anterior midgut and thus exposed to rapid changes in temperature following consumption of a blood meal. DnaK has also been implicated in protection against other environmental stresses, including acidic conditions, antibiotic resistance and oxidative stress (50–52). Our results suggest that *Sodalis* requires DnaK to survive growth *in vitro*, as our attempts to generate mutations in the gene caused cell death. We tried insertions at four different locations in *dnaK* using Targetron mutagenesis, as well as deletion of *dnaK* by allelic change. Notably, these mutagenesis techniques have been successfully used to make mutations in numerous other *Sodalis* genes (53–55). As *Sodalis* transitions from a free-living lifestyle to a mutualistic one (33), growth outside the tsetse host (on agar plates or in liquid BHI) may generate low-grade oxidative stress that requires DnaK for an appropriate cellular response that is critical for bacterial survival.

Maintenance of thermal tolerance homeostasis is an integral process that underlies successful insect host-bacterial symbiont interactions. In fact, symbiotic relationships are disrupted at elevated temperatures, and in some cases, heat-shock can result in the complete loss of these bacteria (reviewed in (56)). Symbionts that live within obligate hematophagous arthropods experience rapid changes in temperature as their host feeds, and these changes must be quickly mitigated to avoid disruption of epidemiologically relevant physiologies. For example, symbiotic bacteria that reside within arthropod vectors indirectly and directly mediate their host’s vector competency. Tsetse’s obligate symbiont *Wigglesworthia* (57), and mosquito (58) and tick (59) commensals, are responsible for mediating production of their host’s peritrophic matrix (PM). This structure is a chitinous and proteinaceous ‘sleeve’ that lines the arthropod midgut (60), and in each of the abovementioned vectors, the PM serves as a protective barrier that their respective vertebrate pathogens must circumvent in order to establish midgut infections that are required for transmission to a subsequent vertebrate host. Intriguingly, *Sodalis* produces chitinase (15) that may exert a two-fold impact trypanosome infection establishment. First, *Sodalis* chitinolytic activity would likely degrade the structural integrity of tsetse’s PM, thus making it easier for trypanosomes to cross the barrier and establish an infection in the fly’s ectoperitrophic space (61, 62). This process would result in the accumulation of *N*-acetyl-D-glucosamine, which would further facilitate trypanosome infection establishment by inhibiting that activity of anti-parasitic tsetse lectins (15). While these theories have never been experimentally proven, they are correlatively validated by the fact that *Sodalis* density positively correlates with trypanosome infection prevalence (16, 19, 63). Finally, symbiotic bacteria from the genera *Kosakonia* and *Chromobacterium*, which are found naturally in the midgut of *Anopheles gambiae* and *Aedes aegypti* mosquitoes, produce and secrete reactive oxygen intermediates (64), histone deacetylases (65) and aminopeptidases (66) that exert direct anti-*Plasmodium* and anti-dengue activity.

In conclusion, information about *Sodalis’* heat shock response provides insight into bacterial adaptations that allow symbionts residing within the gut of hematophagous arthropods to survive acute environmental stressors, including heat shock that ensues immediately after their host consumes a meal of vertebrate blood. This information increases our understanding of the physiological mechanisms that facilitate maintenance of bacterial symbioses, which are crucial mediators of host fitness and vector competency.

## ACKNOWLEDGEMENTS

We gratefully thank Dr. Bernd Bukau at University Heidelberg for supplying us with strain MC4100Δ*dnaK*. We also thank the students in the Introduction to Biological Thinking class at the University of Richmond in the Fall of 2010 for help with preliminary data acquisition.

## SUPPORTING INFORMATION CAPTIONS

**Fig. S1. Non-coding regulatory elements for *Sodalis dnaK*, *dnaK*, and *grpE* genes.** (A) Putative *Sodalis* promoters for the polycistronic *dnaK* and *dnaJ* mRNA and for monocistronic *grpE* mRNA are shown, based on homology to their *E. coli* promoters. Start codons are bolded, the Shine-Delgarno sequence is bolded and italicized, and the σ^32^ binding sites are bolded and underlined. (B) The consensus sequence for the σ^32^ binding site for *E. coli* (67). (C) A potential secondary structure of the RNA corresponding to the *dnaK*–*dnaJ* intergenic region, generated using RNAfold from the ViennaRNA package (68, 69).

**Fig. S2. Comparison of *Sodalis* and *E. coli* heat shock chaperone proteins.** Alignment of *Sodalis* DnaK, DnaJ, and GrpE with homologues from *Escherichia coli* MG1665 using Clustal Omega (https://www.ebi.ac.uk/Tools/msa/clustalo/). An asterisk (*) indicates positions that have a single, fully conserved residue. A colon (:) indicates conservation between groups that exhibit strongly similar properties, roughly equivalent to scoring > 0.5 in the Gonnet PAM 250 matrix. A period (.) indicates conservation between groups that exhibit weakly similar properties, roughly equivalent to scoring ≤0.5 and > 0 in the Gonnet PAM 250 matrix. For DnaK, the boxed residues indicate a glycine (G) that interacts with GrpE, a glutamine (Q) that binds the unfolded protein substrate and an alanine (A) that is involved in synergistic activation of ATPase by DnaJ (70–72). The overlined residues indicate DnaK amino acids predicted to interact with Mg-ADP (71, 73, 74). The dashed underline indicates a motif found in DnaK from all gram-negative bacteria that is thought to be essential for ATP-dependent cooperative function with DnaJ and GrpE (75). The threonine (T) with the dot is required for ATPase activity (76). For DnaJ, the bracketed residues are conserved residues in the J-domain that interact with DnaK (77, 78). The underlined residues are zinc-binding motifs that are predicted to bind the unfolded protein substrate (79–81). The G/F region, which may modulate unfolded substrate binding to DnaK, is boxed, and the DIF motifs within this G/F region, which are involved in regulation of chaperone cycling by modulating a step after ATP hydrolysis (82, 83), are overlined.

**Fig. S3. Comparison of DnaK proteins from *E. coli*, *Sodalis glossinidius,* and other insect symbionts.** Alignment of *Sodalis glossinidius* DnaK with homologues from *Escherichia coli* MG1665 and the insect symbionts using Clustal Omaga (https://www.ebi.ac.uk/Tools/msa/clustalo/). The species corresponding to the protein accession numbers are as follows: WP_074011646.1, *Candidatus Sodalis* sp. SoCistrobi; KYP97672.1, *Sodalis*-like endosymbiont of *Proechinophthirus fluctus*; WP_025244843.1, *Candidatus Sodalis pierantonius*; WP_067565807.1, *Candidatus Doolittlea endobia*; WP_067567978.1, *Candidatus Hoaglandella endobia*; WP_014888228.1, secondary endosymbiont of *Ctenarytaina eucalypti*; WP_067497883.1, *Candidatus Gullanella endobia*; WP_067568929.1, *Candidatus Mikella endobia*; AIN47473.1, *Candidatus Baumannia cicadellinicola*; WP_014888738.1; secondary endosymbiont of *Heteropsylla cubana*; WP_083172452.1, secondary endosymbiont of *Trabutina mannipara*; WP_013975497.1, *Candidatus Moranella endobia*. An asterisk (*) indicates positions which have a single, fully conserved residue. A colon (:) indicates conservation between groups of strongly similar properties, roughly equivalent to scoring > 0.5 in the Gonnet PAM 250 matrix. A period (.) indicates conservation between groups of weakly similar properties, roughly equivalent to scoring =< 0.5 and > 0 in the Gonnet PAM 250 matrix. The boxed residues indicate a glycine (G) that interacts with GrpE, a glutamine (Q) that binds the unfolded protein substrate and an alanine (A) that has been shown to be involved in synergistic activation of ATPase by DnaJ. The overlined residues indicate DnaK amino acids predicted to interact with Mg-ADP. The dashed underline indicates a motif found in DnaK from all gram-negative bacteria which is thought to be essential for ATP-dependent cooperative function with DnaJ and GrpE. The threonine (T) with the dot is required for ATPase activity.

